# Structural disconnections caused by white matter hyperintensities in post-stroke spatial neglect

**DOI:** 10.1101/2024.07.31.605749

**Authors:** Lisa Röhrig, Hans-Otto Karnath

**Affiliations:** Center of Neurology, Division of Neuropsychology, Hertie Institute for Clinical Brain Research, University of Tübingen, 72076 Tübingen, Germany

**Keywords:** attention, connectome, disconnection-symptom mapping, leukoaraiosis, network, predictive modeling

## Abstract

White matter hyperintensities (WMH), a common feature of cerebral small vessel disease, affect a wide range of cognitive dysfunctions, including spatial neglect. The latter is a disorder of spatial attention and exploration typically after right hemisphere brain damage. To explore the impact of WMH on neglect-related structural disconnections, the present study investigated the indirectly quantified structural disconnectome induced by either stroke lesion alone, WMH alone, or their combination. Further, we compared different measures of structural disconnections – voxel-wise, pairwise, tract-wise, and parcel-wise – to identify neural correlates and predict acute neglect severity. We observed that WMH-derived disconnections alone were not associated to neglect behavior. However, when combined with disconnections derived from individual stroke lesions, pre-stroke WMH contributed to post-stroke neglect severity by affecting right frontal and subcortical substrates, like the middle frontal gyrus, basal ganglia, thalamus, and the fronto-pontine tract. Predictive modeling demonstrated that voxel-wise disconnection data outperformed other measures of structural disconnection, explaining 42% of the total variance. Compared to using stroke lesion anatomy, prediction performance can be improved by either estimating stroke-based structural disconnections or delineating the combined anatomy of stroke lesion and WMH. We conclude that pre-stroke alterations in the white matter microstructure due to WMH contribute to post-stroke deficits in spatial attention, likely by impairing the integrity of human attention networks.

## Introduction

Aside from neuronal gray matter damage, pathological behavioral phenomena can also be caused by structural and/or functional disconnection of structurally intact gray matter regions (Catani & Ffytche, 2005; Geschwind, 1965). As a result, structural disconnections are associated with a range of post-stroke cognitive impairments (e.g. den Ouden et al., 2019; Pan et al., 2022; Salvalaggio et al., 2020; Talozzi et al., 2023). Such disconnection can result from damage to the same white matter pathway(s) at different (subcortical) locations, i.e. despite the lack of focal lesion overlap (Catani & Mesulam, 2008). In addition to major stroke events, white matter damage can also originate from cerebral small vessel disease. Manifestations of the latter can be small subcortical infarcts, lacunes, microbleeds, perivascular spaces, brain atrophy, and white matter hyperintensities (WMH) (Wardlaw, Smith, Biessels, et al., 2013; Wardlaw, Smith, & Dichgans, 2013). WMH, also known as leukoaraiosis, are frequently observed in brain imaging of the elderly and appear as hyperdense areas on T2-FLAIR images. They are categorized into periventricular WMH (lesions surrounding lateral ventricles) and deep WMH (punctate to coalescing lesions within the deep/subcortical white matter).

While the exact pathophysiology remains under investigation, WMH may arise from diverse pathological mechanisms that result in histological alterations such as myelin and axonal loss (Gouw et al., 2011). WMH burden was shown to be associated with disturbances in cognitive functions in dementia (for a meta-analysis, see Hu et al., 2021), Parkinson’s disease (e.g. Liu et al., 2021), and post-stroke aphasia (e.g. Vadinova et al., 2024; Wilmskoetter et al., 2019), among others. With respect to WMH-related structural disconnectivity, research previously revealed its relation to various cognitive dysfunctions (Langen et al., 2018; Lee et al., 2021; Respino et al., 2019; Yang et al., 2020).

In this context, the present study addresses the most common and debilitating cognitive disorder after right hemisphere damage, namely spatial neglect – a disorder of spatial attention and exploration. Patients with spatial neglect shift their attention towards their ipsilesional, right side of space while ignoring objects and people located on their contralesional, left side, with a persistent eye and head bias to the right (Coelho-Marques et al., 2023; Fruhmann-Berger & Karnath, 2005). Spatial attention and exploration is proposed to be processed within a perisylvian brain network (Corbetta & Shulman, 2011; Karnath, 2009; Karnath & Rorden, 2012), which is supported by commonly damaged white matter pathways (Bartolomeo et al., 2007; Karnath et al., 2009; Thiebaut de Schotten et al., 2014) and stroke lesion-derived structural disconnections (Saxena et al., 2022; Vaessen et al., 2016; Wiesen et al., 2020). Regarding the impact of WMH on post-stroke spatial neglect, previous studies indicate that a larger extent of WMH correlates with more frequent and severe spatial neglect during both the acute (Bahrainwala et al., 2014; Röhrig et al., 2022) and chronic (Hawe et al., 2018; Kamakura et al., 2017) phase of stroke. This prompts the question of whether alterations in the structural connectome, particularly premorbid changes in the white matter integrity between (sub)cortical areas involved in spatial attention, might contribute to the observed link between WMH extent and severity of post-stroke spatial neglect.

We here investigated the potential influence of WMH on spatial neglect by comparing structural disconnections caused solely by stroke lesions with those resulting from WMH alone or from the individual combinations of stroke lesion and WMH. In addition, we aimed to gain a comprehensive understanding of neglect-related white matter damage: *voxel-wise disconnection* reveals the topography of disconnectivity, *pairwise disconnection* the damage between two gray matter regions, *tract-wise disconnection* the damage along each white matter tract, and *parcel-wise disconnection* evaluates the damage for each gray matter region. We further explored which kind of data yields most accurate predictions of spatial neglect severity (among the mentioned measures of structural disconnection, with and without WMH involvement).

## Methods

### Participants

We investigated 103 patients with an acute, first-time right hemisphere stroke. Patients were consecutively admitted to the Center of Neurology of the University of Tübingen. The sample (Table 1) was previously examined in an earlier study (Röhrig et al., 2022). Patients with bilateral lesions, diffuse demarcations, brain tumors, or without available T2-FLAIR scans were excluded; further, patients with time periods of more than 16 days between stroke onset and brain imaging or neuropsychological assessment were also not included. All patients gave their informed consent for study participation and scientific data usage. The study was approved by the ethics committee of the medical faculty of Tübingen University and was conducted in accordance with the revised Declaration of Helsinki.

**Table 1.** Patient sample (*N*=103).

| Parameter | Neglect (N = 27) | No neglect (N = 76) | p |
| --- | --- | --- | --- |
| Age [years] | 60.5 (15.1) | 57.4 (13.2) |  |
| Sex (F, M) | 10, 17 | 27, 49 |  |
| Stroke onset to imaging [days] | 4.7 (4.7) | 3.1 (3.5) |  |
| Etiology (I, H) | 26, 1 | 70, 6 |  |
| Lesion volume [cm<sup>3</sup>] | 55.6 (44.4) | 17.5 (20.1) | *** |
| WMH volume [cm<sup>3</sup>] |  |  |  |
| ... bilateral | 9.6 (13.0) | 7.5 (6.9) |  |
| ... right hemisphere | 4.5 (6.0) | 3.7 (3.4) |  |
| Stroke onset to assessment [days] | 3.0 (2.3) | 3.5 (3.0) |  |
| Letter CoC | 0.39 (0.30) | 0.01 (0.02) | *** |
| Bells CoC | 0.46 (0.27) | 0.02 (0.03) | *** |
| Mean CoC | 0.43 (0.28) | 0.01 (0.02) | *** |
| Visual field defects [N] | 3 | 9 |  |

*Note.* Patients with mean CoC $\geq 0.082$ were defined as having spatial neglect (cut offs for the letter and bells cancellation tasks are 0.083 and 0.081, respectively; Rorden & Karnath, 2010). Results are given in either mean (standard deviation) or number of patients. “*Lesion/WMH volume*” represents the normalized volume in MNI space. Significant group differences are highlighted with asterisks (\*\*\* – $p < 0.001$ ).
*Abbreviations:* F – females, M – males, I – Ischemia, H – Hemorrhage.

### Behavior

We used the standard paper-pencil tests letter cancellation (Weintraub & Mesulam, 1985) and bells cancellation (Gauthier et al., 1989) to test patients for spatial neglect. A continuous score depicting the severity of neglect was calculated: the Center of Cancellation (CoC; Rorden & Karnath, 2010) captures both the number and localization of omissions and ranges between-1 (right-sided neglect) and +1 (left-sided neglect). The mean CoC was calculated for each patient that was used for the analyses (in five patients, one of both CoC-scores was missing, thus, the available CoC-score was used). As the CoC is a bipolar variable, where CoC = 0 represents the optimum and positive and negative values represent spatial biases in attention, we set negative mean CoC-values to 0 (right-sided neglect; *N* = 13, all not pathological) in order to increase interpretability and to prevent that the algorithm assumes negative scores as less impaired than scores ≈ 0. Visual field defects were assessed by the standard neurological confrontation technique.

### Imaging

MR images including a T2-FLAIR were available from all patients. Stroke lesion maps were delineated on DWI (diffusion-weighted imaging) if imaging was acquired within 48 hours post-stroke (*N* = 47) or on T2-FLAIR scans (*N* = 56). WMH were delineated on (co-registered) T2-FLAIR images. Both delineation procedures were carried out using the semi-automated “Clusterize Toolbox” (de Haan et al., 2015) for SPM12 (Wellcome Department of Imaging Neuroscience, London, UK) for MATLAB R2019a (The MathWorks, Inc., Natick, USA). Stroke lesion maps were then normalized to MNI (Montreal Neurological Institute) space with a voxel size of 1×1×1 mm using age-matched templates distributed by the “Clinical Toolbox” (Rorden et al., 2012); WMH maps were normalized by applying the same transformation parameters as for the patients’ stroke lesion maps. The final bilateral WMH maps were separated at the midsagittal plane to obtain right hemispheric WMH maps. Further details on the imaging preprocessing steps can be found in Röhrig et al. (2022).

Patients’ voxel-based binary lesion maps were used to indirectly obtain structural disconnection estimates. For this purpose, we used the “Lesion Quantification Toolkit” (LQT; Griffis et al., 2021). It allows to indirectly estimate white matter disconnections in brain-damaged patients by implementing the individual lesion map as a “seed region”: streamlines of the normative connectome that intersect with a lesion are estimated to be affected (Griffis et al., 2021; Sperber et al., 2022). We used the default DTI white matter tractography atlas of 842 healthy individuals of the Human Connectome Project (HCP-842; Yeh et al., 2018) and the “Brainnetome Atlas” (BNA) as gray matter parcellation that is based on structural and functional connectivity measures (Fan et al., 2016). The BNA comprises 246 (i.e. 210 cortical and 36 subcortical) subregions of bilateral hemispheres. The LQT was set to consider only end-wise connections (i.e. direct connections that end in both gray matter regions, rather than just passing through them), as recommended for interpretability reasons.

With respect to structural disconnection metrics, we used relative disconnection values (ranging between 0 and 100 percent) in either individual (1) 3D voxel-wise disconnection maps, (2) 2D symmetric matrices of pairwise (region-to-region, connection-wise) disconnections, (3) 1D tract-wise disconnections, or (4) 1D parcel-wise disconnections. With respect to structural disconnection origins, we tested the contribution of WMH by analyzing (i) stroke lesion-induced disconnections, (ii) WMH-induced disconnections, and (iii) disconnections induced by the combination of stroke lesion and WMH. As we revealed in a previous investigation that, first, right hemispheric and bilateral WMH outperformed left hemispheric WMH and, second, the topography of right hemispheric and bilateral WMH were almost equally predictive (Röhrig et al., 2022), we focus on right-sided WMH in the following. However, we also present results from equivalent analyses using bilateral WMH in the Supplement.

### Data analyses

For all subsequent analyses, we considered one-sided *p*-values of maximum 0.05 as significant. Our analyses aimed to find associations with positive statistics between white matter disconnection and neglect severity.

### Voxel-wise analyses

To investigate, whether specific voxels of the white matter disconnectome are related to spatial neglect, we employed mass-univariate voxel-based disconnectome-symptom mapping (VDSM) by implementing general linear models (GLMs) using NiiStat (https://www.nitrc.org/projects/niistat). The approach is similar to the well-known voxel-based lesion-symptom mapping (VLSM) but relies on structural disconnection maps instead of lesion anatomy maps. Continuous disconnection maps were binarized at a threshold of > 50 % disconnection (Wawrzyniak et al., 2022; Wiesen et al., 2020). The mean CoC-scores served as the outcome measure. To reduce the amount of data with zero to low variability, we excluded voxels that were damaged in less than 5 patients as it is a common practice in voxel-based analyses (e.g. Wawrzyniak et al., 2022). Analysis-specific numbers of investigated features are reported in Table S1 in the Supplement. To correct for multiple testing, we used the permutation-based family-wise error (FWE) correction with 10,000 iterations. To identify voxels that were more strongly associated to neglect severity when WMH were considered, we calculated *z*-statistics obtained by stroke lesion and WMH minus *z*-statistics obtained by stroke lesion only (*Δz*); we only report voxels that were tested in both analyses, that resulted in *Δz* > 0.01, and that reached significance when WMH were considered (*p* < 0.05).

### Pairwise analyses

As a second step, we employed connectome-based lesion-symptom mapping (CLSM) to explore alterations at the network-level (Gleichgerrcht et al., 2017). We computed pairwise (region-to-region, connection-wise) analyses with custom scripts in MATLAB R2019a that were used similarly in a previous study (Röhrig et al., 2023). Hereby, we investigated whether damage to direct structural connections between specific brain regions are associated with the severity of acute post-stroke neglect. For this procedure, we used the symmetric pairwise matrices obtained from the LQT and deleted redundant cells. Similar to the voxel-wise mapping, we excluded features with low variability to avoid less meaningful results and a very large number of features to be tested. Pairwise analyses were restricted to connections that were damaged in at least 10 patients (Tab. S1). Like the voxel-wise analyses, we implemented mass-univariate GLMs to identify connections associated with the behavioral deficit. The maximum *t*-statistics approach based on 10,000 permutations was used to correct for multiple testing. To identify pairwise connections that were more strongly associated to neglect severity when WMH were considered, we calculated *t*-statistics obtained by stroke lesion and WMH minus *t*-statistics obtained by stroke lesion only (*Δt*), as for the voxel-wise analysis.

### Tract-wise analyses

Tract-wise analyses were carried out in the same way as the analyses using pairwise disconnections, to identify white matter tracts associated to neglect severity. We used the vector with tract-level disconnection severities derived from the LQT that contains the proportion of streamlines of each white matter tract that intersect with the lesion. The LQT uses 66 canonical tracts of the HCP-842 tractography atlas (Yeh et al., 2018) while splitting the corpus callosum into 5 compartments based on the FreeSurferSeg ROIs distributed with DSI Studio (https://dsi-studio.labsolver.org/) (see Griffis et al., 2021). From the total of 70 tracts, we excluded those that were damaged in less than 10 patients (Tab. S1).

### Parcel-wise analyses

Parcel-wise analyses were carried out in the same way as pairwise analyses, to investigate the influence of accumulated damage to single gray matter regions on neglect severity. We used the vector with parcel-level disconnection severities derived from the LQT that contains the percentage of parcel damage. As we used the BN-atlas, 246 regions were analyzed. Analyses were restricted to parcels that were damaged in at least 10 patients (Tab. S1).

### Predictive modeling

Lastly, we implemented predictive modeling to investigate the predictive power of voxel-wise, pairwise, tract-wise, and parcel-wise structural disconnections derived from either stroke lesions/WMH alone or the combination of stroke lesion and WMH to predict acute severity of spatial neglect. We applied the supervised learning algorithm support vector regression (SVR) with a nonlinear radial basis function kernel and a repeated nested cross-validation procedure to prevent overfitting (i.e. 10-fold outer loop and 5-fold inner loop, with 10 model repetitions). Mean CoC-scores were square-root-transformed and served as the test score. We ignored features that were less frequently damaged across patients (see above). Dimensions of continuous voxel-wise measures were reduced via principal component analysis (see legend of Tab. S1). The algorithm was designed to minimize the mean squared error (MSE). The coefficient of determination (R²) was used to estimate model’s goodness of fit; it gives the proportion of variance explained by the model. For further details on the prediction algorithm, see Röhrig et al. (2022).

## Results

Overlay plots of stroke lesions and WMH as well as related white matter disconnection topographies are illustrated for patients with spatial neglect (*N* = 27) and without neglect (*N* = 76) in Figure 1. For the reasons mentioned above, results of analyses relating to right-sided WMH are presented in the following. Results of equivalent analyses with bilateral WMH are reported in the Supplement. Overall, findings were relatively similar for both WMH variants, although analyses using right-sided WMH generally found a greater number of significant associations. With respect to analyses using only WMH-induced structural disconnection data, we found no voxels, pairwise connections, tracts, or parcels significantly associated to neglect severity (for both right-sided and bilateral WMH; *p* > 0.05). This indicates that WMH burden alone (without a stroke) is not linked to neglect behavior. In addition, WMH-derived disconnection data were not predictive for neglect severity (*R²* ≈ 0).

**Figure 1.**
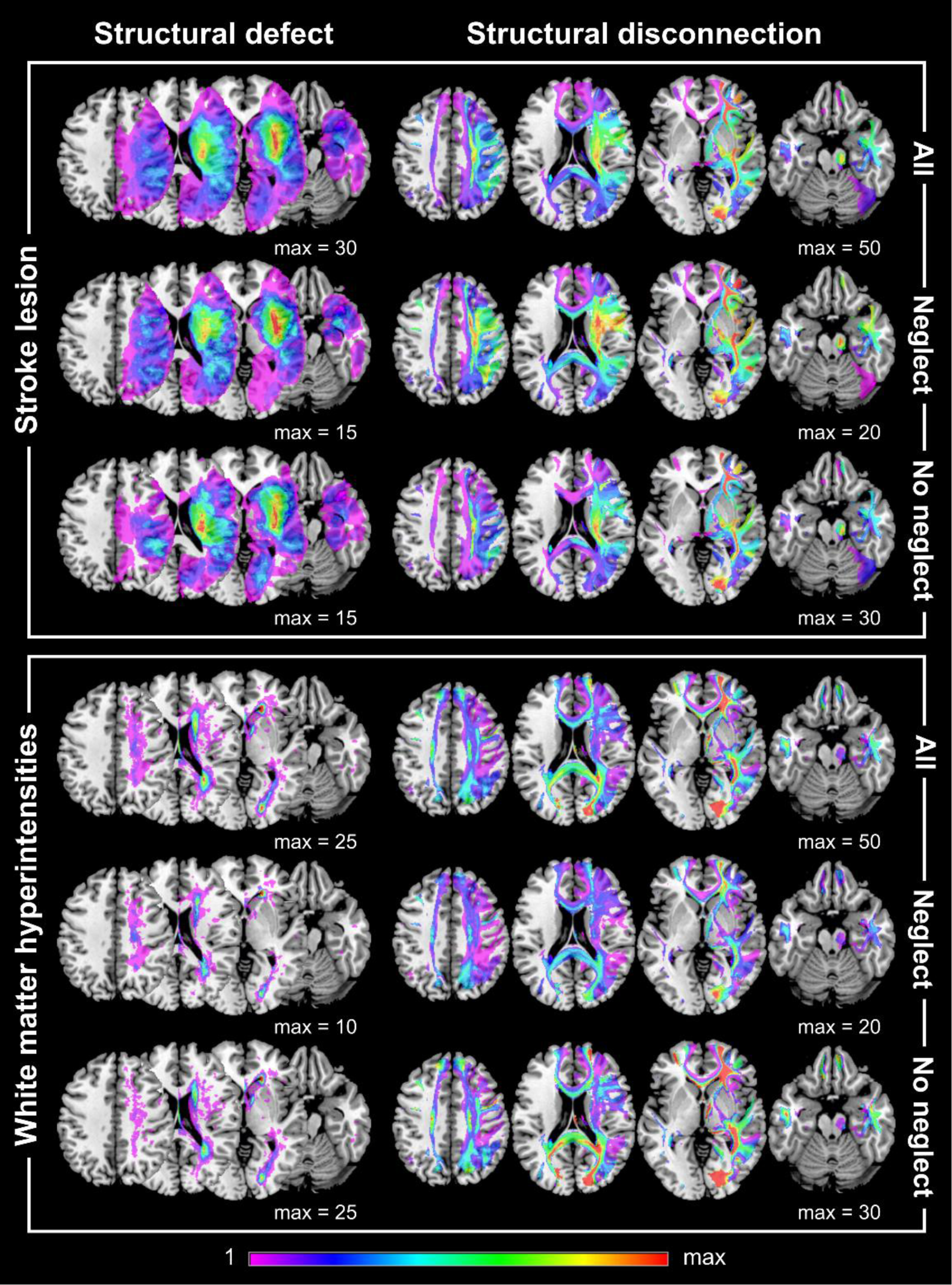
Overlay plots. Overlay plots of anatomical lesion maps and corresponding structural white matter disconnection maps are presented for right-sided stroke lesions (upper panel) and right-sided WMH (lower panel) on a brain template in MNI space. Topographic maps are shown for the total sample (*N* = 103, upper row each) and for patients with spatial neglect (*N* = 27, middle row each) and without (*N* = 76, lower row each). Structural disconnection maps were binarized (> 50 % disconnection). Brain slices with *z*-coordinates of 40, 20, 0, and-20 are presented. The color bar depicts the number of patients who have damage to a specific voxel; “*max”* represents the maximum value used for the corresponding color map.

### Voxel-wise structural disconnections

As a first step, we computed whole-brain analyses to investigate whether specific voxels of the disconnection topography are associated with neglect severity. With respect to stroke lesion-induced structural disconnections, one patient had to be excluded as the corresponding disconnection map did not contain any voxels with more than 50 % disconnection. 4885 voxels derived from the stroke lesion-induced disconnectome were found to be significantly associated with neglect severity (*z* > 5.33; Fig. 2A). The voxel with the maximum *z*-score of 6.95 is part of the right ventrolateral area 8 within the middle frontal gyrus (MFG) (MNI-coordinates = 34, 20, 52). Among white matter tracts, significant voxels were most frequently part of the corpus callosum, right cortico-striatal pathway, and right cortico-thalamic pathway (HCP-842; Yeh et al., 2018). Moreover, significant voxels cover also white matter underlying the MFG, superior temporal gyrus (STG), and inferior parietal lobule (IPL) (Brainnetome Atlas; Fan et al., 2016).

**Figure 2.**
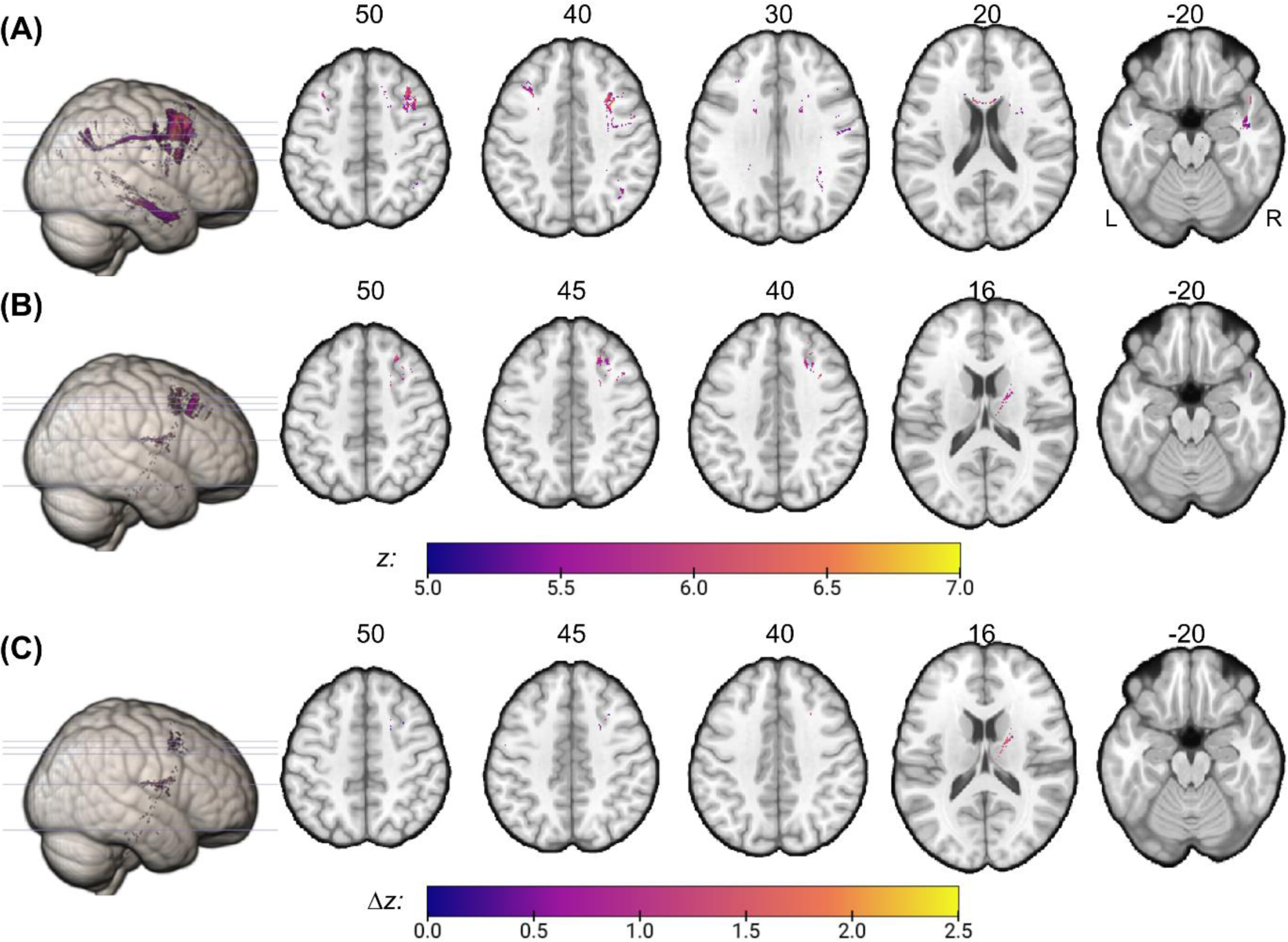
Voxel-wise results. Voxels of the structural disconnectome are colored that were found to be significantly associated with neglect severity (*p* < 0.05, with permutation-based FWE-correction). Structural disconnection maps were binarized at 50 % disconnection and were derived from either **(A)** the stroke lesion alone or **(B)** the combination of stroke lesion and right-sided white matter hyperintensities (WMH). Voxel masks are presented on a ch2-template (numbers above axial slices refer to the corresponding z-coordinate in standard MNI space). The color-bar refers to the obtained statistical *z*-scores. **(C)** shows results obtained by subtracting **(A)** from **(B)**; Δ*z*-values demonstrate increased strength of association between disconnection severity and neglect severity due to right-sided WMH.

When analyzing the combined topography of stroke lesion and right hemispheric WMH, we identified 937 voxels that survived FWE-correction (*z* > 5.59; Fig. 2B). The voxel with the maximum *z*-score of 6.84 is part of the right inferior frontal junction (IFJ) within the MFG (MNI-coordinates = 39, 8, 47). Among white matter tracts, significant voxels were most frequently part of the right fronto-pontine tract, cortico-thalamic pathway, and cortico-striatal pathway. Significant voxels further cover white matter underlying the superior frontal gyrus, MFG, and thalamus. In comparison to stroke-derived results that nicely reveal areas of the perisylvian network including frontal, parietal, and temporal regions (Fig. 2A), the analysis that considers WMH reveals a different distribution of significant voxels, namely predominantly fronto-subcortical locations (Fig. 2B). A similar voxel distribution is also demonstrated by the subtraction of obtained statistics that resulted in 300 voxels with a stronger association to neglect behavior when right-sided WMH were considered (Fig. 2C); the voxel with the largest increase is located at the right cortico-thalamic pathway next to the dorsal caudate (MNI-coordinates = 20, 2, 20).

### Pairwise structural disconnections

We further implemented pairwise (region-to-region) analyses to investigate the association of neglect severity and the proportional damage to direct structural white matter connections between two regions. We found significant pairwise disconnections after permutation-based FWE-correction. Stroke lesion-induced disconnectivity revealed the following number of significant disconnections: *N* = 394 at *p* < 0.05 (*t* ≥ 4.40; Fig. 3A); *N* = 79 at *p* < 0.01 (*t* ≥ 5.26); *N* = 43 at *p* < 0.005 (*t* ≥ 5.60; Fig. 3A); *N* = 2 at *p* < 0.001 (*t* ≥ 6.64). In total, 143 gray matter regions were involved. Regions with at least 10 disconnections were part of the right superior, middle, and inferior gyri of temporal and frontal lobules, precentral gyrus, insular gyrus, basal ganglia (globus pallidus, putamen) and IPL, and the left middle temporal gyrus (MTG) and IPL. The most significant disconnections were found between left inferior frontal gyrus (IFG) (opercular area 44) and right MFG (ventrolateral areas 6 respectively 8).

**Figure 3.**
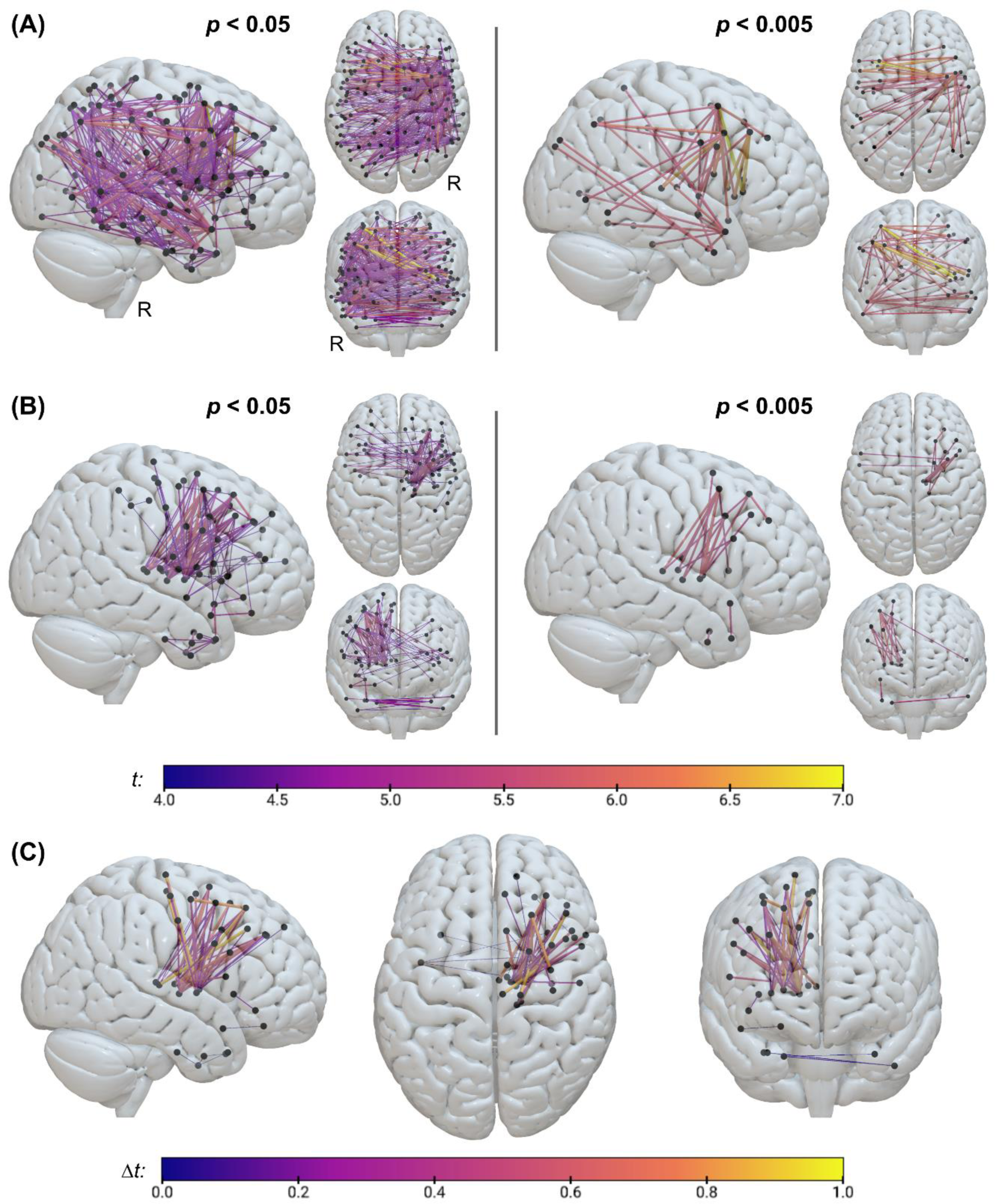
Pairwise results. The plot shows pairwise structural disconnections associated with spatial neglect severity, which were derived from either **(A)** the stroke lesion alone or **(B)** the joint topography of stroke lesion and right-sided WMH. Depicted disconnections are significant at *p* < 0.05 (left) respectively *p* < 0.005 (right), after surviving permutation-based FWE-correction. Nodes are based on the Brainnetome atlas with 246 regions (Fan et al., 2016). Nodes are presented in black, whereas the edge colors represent *t*-values obtained from the respective analysis (see color bar). **(C)** shows results obtained by subtracting **(A)** from **(B)**; Δ*t*-values demonstrate increased strength of association between disconnection severity and neglect severity due to right-sided WMH. R – right hemisphere.

When combining stroke lesion and right-sided WMH, we found the following number of significant disconnections: *N* = 112 at *p* < 0.05 (*t* ≥ 4.21; Fig. 3B); *N* = 41 at *p* < 0.01 (*t* ≥ 4.86); *N* = 18 at *p* < 0.005 (*t* ≥ 5.25; Fig. 3B); *N* = 2 at *p* < 0.001 (*t* ≥ 5.91). In total, 63 gray matter regions were affected. Regions with at least 10 disconnections were part of the right basal ganglia (globus pallidus, putamen), middle and superior frontal gyri, and thalamus. The most significant disconnections were found between right globus pallidus and right MFG (IFJ respectively ventrolateral area 6). Like the voxel-wise results, pairwise findings of stroke lesion-derived disconnections involve frontal, parietal and temporal regions (Fig. 3A), whereas stroke and WMH-derived disconnections involve rather frontal and subcortical areas (Fig. 3B). A similar image is also demonstrated by the subtraction of obtained statistics that resulted in 56 connections with a stronger association to neglect behavior when right-sided WMH were involved (Fig. 3C); the pairwise connection with the largest increase connects the right ventrolateral area 8 of the MFG and the right posterior parietal thalamus. Table S2 in the Supplement reports most significant pairwise disconnections; Table S3 and Figure S1 present most frequently affected regions; Table S6 reports connections with the largest increase in association strength due to WMH.

### Tract-wise structural disconnections

With respect to stroke lesion-based tract-wise analyses, we identified white matter tracts significantly associated to neglect severity: *N* = 22 at *p* < 0.05 (*t* > 3.26); *N* = 17 at *p* < 0.01 (*t* > 4.05); *N* = 13 at *p* < 0.005 (*t* > 4.34); *N* = 6 at *p* < 0.001 (*t* > 5.13; Fig. 4A). The most significant tracts include central and mid-anterior parts of the corpus callosum, right cortico-striatal pathway, right cortico-thalamic pathway, right SLF, and right U-fibers (Fig. 4A/D).

**Figure 4.**
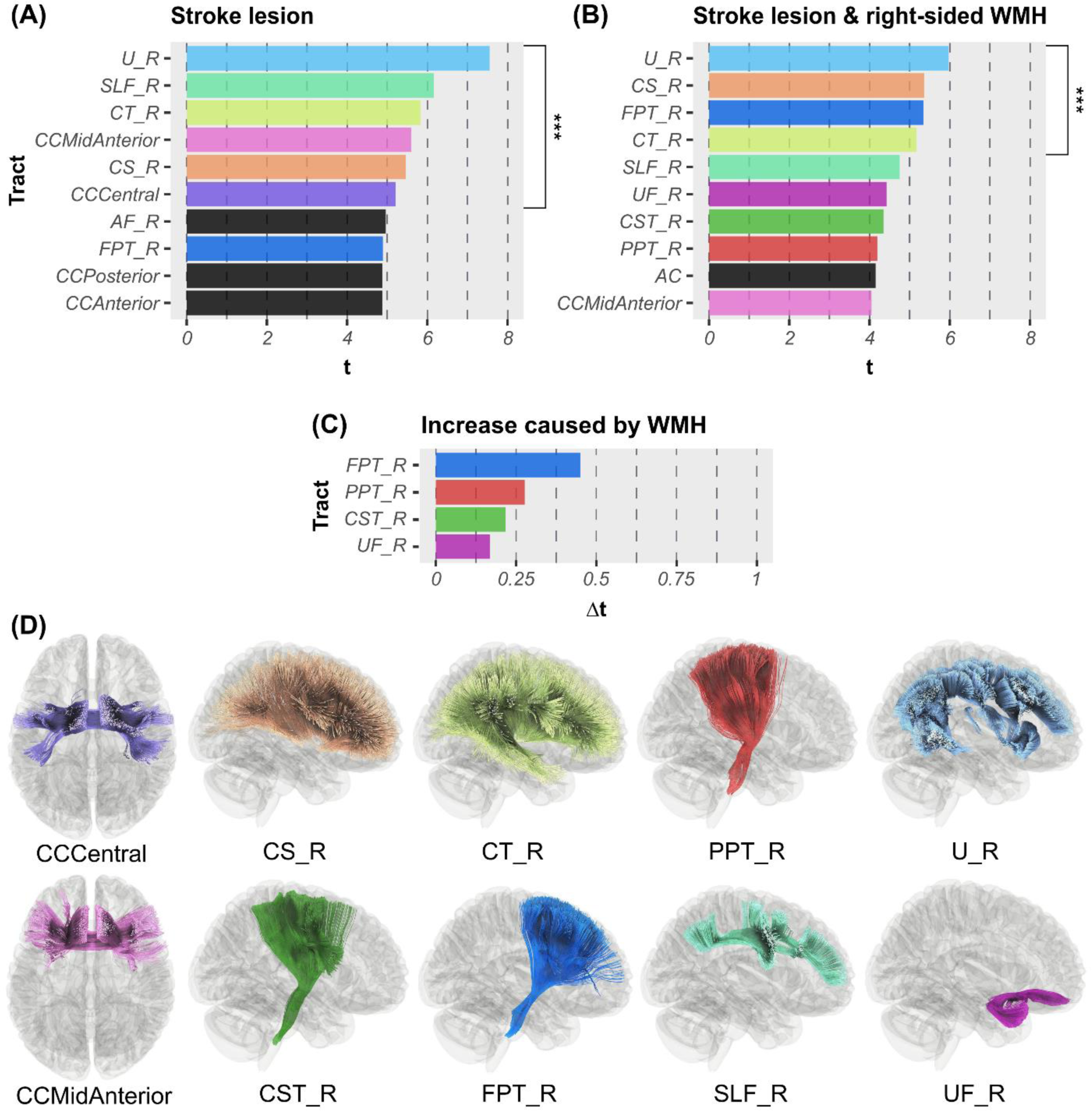
Tract-wise results. **(A/B)** *t*-statistics are presented for the 10 most significant associations between tract-level disconnection severities and spatial neglect severity, revealed by mass-univariate GLMs with permutation-based FWE-correction. Associations were significant **(A)** for stroke lesion-induced disconnections and/or **(B)** for disconnections induced by the combination of stroke lesion and right-sided WMH. Tracts significant at *p* < 0.001 are highlighted (***). **(C)** shows results obtained by subtracting **(A)** from **(B)**; Δ*t*-values demonstrate increased strength of association between disconnection severity and neglect severity due to right-sided WMH. **(D)** White matter tracts based on the HCP-842 atlas (Yeh et al., 2018) are depicted within a glass-brain. *Abbreviations*: AC – anterior commissure; AF – arcuate fasciculus; CC – corpus callosum; CS – cortico-striatal pathway; CST – cortico-spinal tract; CT – cortico-thalamic pathway; FPT – fronto-pontine tract; PPT – parieto-pontine tract; R – right hemispheric; SLF – superior longitudinal fasciculus; U – U-fiber; UF – uncinate fasciculus.

The joint topography of stroke lesion and right-sided WMH did reveal tracts significantly related to neglect severity: *N* = 14 at *p* < 0.05 (*t* > 3.12); *N* = 11 at *p* < 0.01 (*t* > 3.91); *N* = 8 at *p* < 0.005 (*t* > 4.17); *N* = 4 at *p* < 0.001 (*t* > 4.81; Fig. 4B). The most significant tracts include, as for the lesion-based analysis, right cortico-striatal pathway, cortico-thalamic pathway, and U-fibers; in addition, the right fronto-pontine tract was found to be highly associated to neglect severity when right-sided WMH were combined with the stroke lesion (Fig. 4B/D). However, associations were generally less strong for analyses with WMH compared to analyses without WMH. The subtraction of obtained statistics resulted in 4 white matter tracts with a stronger association to neglect behavior when right-sided WMH were considered (Fig. 4C/D); the right fronto-pontine tract received the largest increase in association strength. Table S4 in the Supplement reports significant tract-wise results; Table S6 reports tracts with the largest increase in association strength due to WMH.

### Parcel-wise structural disconnections

Parcel-wise analyses of stroke lesion-derived structural disconnectivity revealed the following significant associations of parcel damage with neglect severity: *N* = 13 at *p* < 0.05 (*t* > 4.38); *N* = 5 at *p* < 0.01 (*t* > 5.25; Fig. 5A); *N* = 3 at *p* < 0.005 (*t* > 5.60). Most significant parcels were right rostral area 21 of the MTG, medial area 38 of the STG, and caudal ventrolateral area 6 of the precentral gyrus (Fig. 5A/C).

**Figure 5.**
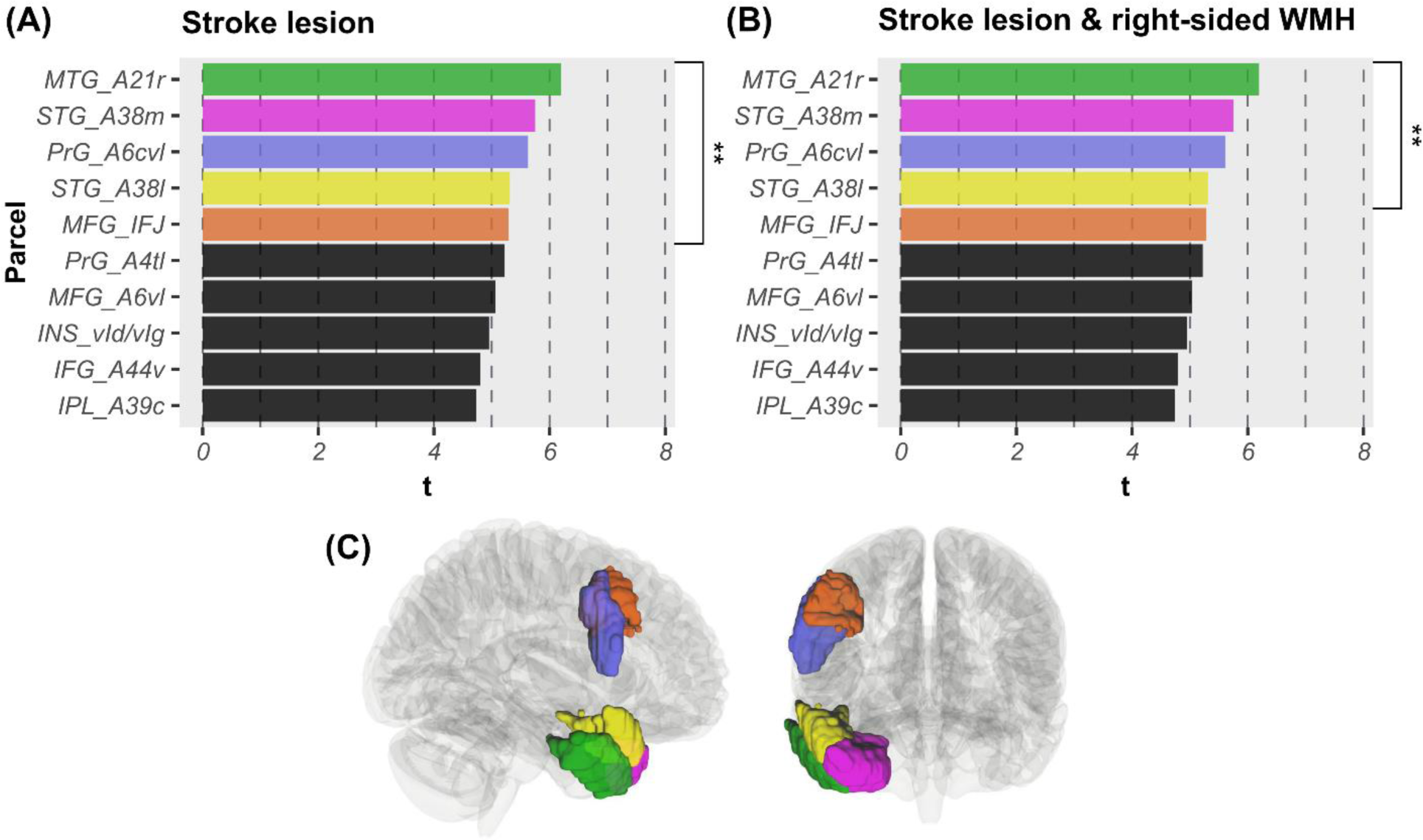
Parcel-wise results. **(A/B)** *t*-statistics are presented for the 10 most significant associations between parcel damage and spatial neglect severity, revealed by mass-univariate GLMs with permutation-based FWE-correction. Associations were significant **(A)** for stroke lesion-induced disconnections and/or **(B)** for disconnections induced by the combination of stroke lesion and right-sided WMH. Parcels significant at *p* < 0.01 are highlighted (**) and visualized in **(C)** with color coding; all significant parcels are right hemispheric. **(C)** Parcels based on the gray matter Brainnetome Atlas (Fan et al., 2016) are displayed within a glass brain. *Abbreviations*: IFG – inferior frontal gyrus; INS – insular gyrus; IPL – inferior parietal lobule; MFG – middle frontal gyrus; MTG – middle temporal gyrus; PrG – precentral gyrus; STG – superior temporal gyrus.

When right-sided WMH were combined with the individual stroke lesion, findings revealed gray matter parcels significantly related to neglect severity: *N* = 10 at *p* < 0.05 (*t* > 4.53); *N* = 4 at *p* < 0.01 (*t* > 5.29; Fig. 5B); *N* = 2 at *p* < 0.005 (*t* > 5.61). Most significant parcels were rostral area 21 of the MTG and medial area 38 of the STG of the right hemisphere (Fig. 5B/C). Note that results for both conditions (i.e. stroke lesions with and without WMH) were very similar with only slightly different values, indicating that WMH mostly affect other parcels than those affected by stroke lesions. Accordingly, the subtraction of obtained statistics demonstrated that no parcel was stronger associated to neglect behavior when WMH were considered. Table S5 and Figure S2 in the Supplement present significant parcel-wise results.

### Prediction analyses

Results obtained by predictive modeling are reported in Table 2. Overall, voxel-wise measures were most predictive and parcel-wise measures least predictive, indicating that high-dimensional data outperformed low-dimensional measures. Most accurate predictions were achieved by voxel-wise structural disconnections derived from individual stroke lesions: almost 42% of the total behavioral variance were explained. For voxel-wise and pairwise measures, stroke lesion-derived disconnectivity outperformed disconnectivity derived from the combination of stroke lesion and WMH. However, bilateral WMH seem to increase the performance for tract-wise data when combined with the stroke lesion, whereas right-sided WMH did slightly increase prediction accuracy for parcel-wise data. Bilateral WMH were more predictive than right-sided WMH when combined with the stroke lesion, except for the analysis using parcel-wise data.

**Table 2.** Prediction performances (*R²*).

| Disconnection origin | Disconnection type |  |  |  |
| --- | --- | --- | --- | --- |
|  | Voxel-wise | Pairwise | Tract-wise | Parcel-wise |
| Stroke lesion | **0.419** | **0.396** | 0.277 | 0.241 |
| Stroke lesion + right-sided WMH | 0.327 | 0.206 | 0.245 | **0.253** |
| Stroke lesion + bilateral WMH | 0.384 | 0.359 | **0.328** | 0.239 |

*Note.* Performances of models predicting acute neglect severity are reported. Predictor variables were measures of structural disconnection derived indirectly either from the stroke lesion or the combination of stroke lesion and right-sided/bilateral WMH. The coefficient of determination ( $R^2$ ) served as the variable that estimates goodness of fit; it gives the proportion of variance explained by a model. Bold values highlight the most accurate performance for each type of disconnection.

## Discussion

The present study investigated whether the combination of stroke lesion and white matter hyperintensities (WMH) lead to more severe structural disconnections between attention-related areas, compared to analyses without WMH involvement. Voxel-wise, pairwise, and tract-wise results were largely consistent, indicating that WMH affect the right frontal and subcortical brain, while WMH do not seem to alter disconnections at the parcel-level. In particular, we observed that WMH, when combined with the individual stroke lesion, predominantly affected connectivity related to the right MFG, basal ganglia and thalamus, as well as the right fronto-pontine tract. Damage to these regions and tracts seemed to be more related to neglect severity when WMH were considered, compared to analyses using only stroke lesion-derived information. Findings therefore suggest that pre-stroke alterations of white matter integrity due to WMH have an impact on deficits in spatial attention following stroke by affecting the neglect-related structural disconnectome. On the other hand, structural disconnections derived from WMH alone were not associated to neglect behavior. Although WMH affect the disconnectome, the individual statistical associations were generally less strong when WMH were involved. This indicates that WMH add noise to the analyses, possibly because not all WMH are similarly related to neglect pathology.

### Neural correlates affected by WMH

A recent study investigating stroke lesion-induced pairwise structural disconnections identified the right superior parietal lobule, insula, and MTG with temporal pole as major hubs impaired in spatial neglect (Wiesen et al., 2022). A similar study by Saxena et al. (2022) found that pairwise disconnections involving the right putamen or right frontal areas were linked to patients with only egocentric neglect behavior. The same study demonstrated critical disconnections involving the right IPL, orbitofrontal cortex, and thalamus when comparing patients with egocentric and/or allocentric neglect versus patients without neglect. While present results of stroke lesion-derived disconnections showed similar findings, present analyses of the combined damage by stroke and WMH revealed a different image of frontal and subcortical regions and tracts being most strongly associated to neglect severity. The association strength of frontal and subcortical disconnections was increased by WMH, suggesting that WMH not only alter the structural connectome but also worsen deficits in spatial attention and exploration post-stroke.

Present findings implicate that WMH contribute to voxel-wise and pairwise disconnection of areas within the right MFG, especially ventrolateral areas 8 and 6 and the inferior frontal junction (IFJ). The right MFG was frequently associated to neglect pathology (Chechlacz et al., 2012; Committeri et al., 2007; Thiebaut de Schotten et al., 2014; for a meta-analysis on neglect anatomy, see Moore et al., 2023). Research shows that lesions in the ventral frontal cortex cause more severe neglect symptoms than temporo-parietal lesions (Rengachary et al., 2011). In accordance, a previous study found a link between neglect and disconnection of the IFJ (Griffis et al., 2021). A meta-analysis demonstrated that the IFJ is functionally co-activated with frontal, temporal, parietal and subcortical regions (Sundermann & Pfleiderer, 2012), of which some were linked to spatial neglect in the present study.

Moreover, we found pairwise structural disconnections between areas of the MFG and basal ganglia when considering right-sided WMH. Basal ganglia were previously linked to neglect symptoms (Chechlacz et al., 2012; Karnath et al., 2002, 2004). It was shown that neglect patients with basal ganglia infarction demonstrate disturbed perfusion in remote cortical areas as the STG, MTG, IPL and IFG (Karnath et al., 2005). As the STG is connected with the putamen and caudate nucleus, it was suggested that a cortico-subcortical network might be involved in spatial awareness (Karnath, 2001). Beyond basal ganglia, present findings also suggest that WMH alter thalamic connections. Thalamic lesions were previously associated to neglect pathology (Karnath et al., 2002; Ten Brink et al., 2019). Nevertheless, studies on perfusion data implied that subcortical infarcts themselves do not cause neglect symptoms, but rather the accompanying cortical hypoperfusion (Hillis et al., 2005; Karnath et al., 2005). Similarly, a very recent investigation on isolated damage of subcortical areas found no strong statistical evidence for an association between isolated basal ganglia damage and spatial neglect (and between isolated thalamus damage and spatial neglect), even though analyses revealed small clusters of functional disconnections between basal ganglia and middle/inferior frontal regions (Sperber et al., 2024).

Frontally located (periventricular) WMH may disrupt pathways between frontal areas and basal ganglia, and between frontal areas and thalamus. Present tract-wise results show that damage to the fronto-pontine tract is particularly more strongly associated to neglect severity due to the involvement of WMH. Beyond, present findings indicate that the parieto-pontine and cortico-spinal tracts are affected by WMH and thereby contribute to worsened neglect. A recent study described a single case of a right pontine lesion causing left-sided spatial neglect (Cazzoli et al., 2023). By using fiber tracking, the authors observed a disruption of bilateral cortico-ponto-cerebellar tracts, which likely include fronto-pontine, parieto-pontine, and cortico-spinal tracts. Disconnection of the cortico-ponto-cerebellar tract was also revealed as a critical difference between patients with and without spatial neglect (Thiebaut de Schotten et al., 2014). Despite very strong associations, WMH have, however, not strengthened the association of cortico-striatal and cortico-thalamic pathways. Contrary to the tract-level, at the voxel-level WMH have most intensively increased the association of the cortico-thalamic pathway. Present results further demonstrate that the uncinate fasciculus was more strongly associated to neglect pathology due to WMH. Damage to this fiber was previously linked to neglect symptoms (Karnath et al., 2011). In fact, the uncinate fasciculus appears to be the white matter tract second most frequently associated with egocentric neglect symptoms in a recent meta-analysis (Moore et al., 2023).

### WMH impact on attention networks

The present study revealed that WMH-derived information influenced voxel-wise, pairwise and tract-wise results, whereas they did almost not influence parcel-wise results. In other words, WMH damage does not concentrate on connectivity of specific (neglect-related) gray matter regions, but rather affects white matter tracts and connections between two gray matter regions that are related to neglect symptoms. This might indicate that WMH impact the integrity of widespread brain networks. Accordingly, previous lesion network mapping showed that visual, ventral attention, and fronto-parietal networks were most susceptible to WMH-induced functional disconnectivity in older participants with cerebral small vessel disease (Crockett et al., 2021). This is supported also by fMRI-based findings, as participants with WMH presented less activity in fronto-temporal and parietal areas compared to participants without WMH (Atwi et al., 2018); the authors further observed absent activation patterns related to processing speed in participants with WMH and concluded that WMH may provoke deficits in attention. In a very recent investigation of a large group of memory clinic patients, WMH-derived structural and functional disconnectivity scores obtained from regions of the dorsal and ventral attention networks were significantly associated to worse performances across cognitive domains (Petersen et al., 2024). In addition, a recent study by Bonkhoff and colleagues (2022) showed that WMH burden is linked to the severity of strokes affecting temporo-parietal regions of the right hemisphere; also, subcortical stroke lesions were related to unfavorable outcomes in that study.

These previous findings highlight that WMH burden influences the impact of strokes involving attention-related regions; they support present findings that structural disconnectivity of frontal areas like the IFJ caused by the combination of stroke and WMH is more strongly linked to neglect behavior compared to when WMH were not considered. The IFJ within the MFG is activated in both dorsal and ventral attention networks (He et al., 2007), suggesting that this region is involved in integrating information between these networks. Altogether, WMH seem to negatively impact post-stroke spatial attention, albeit WMH-related damage alone do not cause spatial neglect. This observation is in accordance with the concept of “brain reserve” (first reviewed in Satz, 1993; details on implications for spatial neglect in Umarova, 2017). It hypothesizes that a particularly healthy brain can tolerate the effects of an extreme event like a stroke due to a sufficiently great reserve, unlike a pre-damaged brain, which would be more vulnerable to stroke-related impacts on brain function. Across studies, WMH were shown to decrease the individual brain reserve and negatively affect brain plasticity, leading to earlier and more severe symptoms after brain damage (Galluzzi et al., 2008).

### The role of structural disconnections in predictive modeling

A further aim of the present study was to explore which kind of data ‒ regarding different measures of structural disconnection, with and without WMH contribution ‒ yields most accurate predictions of severity of spatial neglect. Overall, prediction accuracy was lower for models that included WMH. This observation indicates that WMH-derived disconnections may be noisy and cannot improve the already quite accurate model that uses only stroke lesion-derived disconnection data. Tract-wise data may present an exception, where bilateral WMH seem to clearly benefit model accuracy. Bilateral WMH-derived structural disconnections seem to be generally more predictive than data obtained from right-sided WMH.

We further observed that higher dimensional measures of structural disconnection outperformed lower dimensional measures: voxel-wise structural disconnection was revealed as the most predictive measure, followed by pairwise and tract-wise measures, whereas parcel-wise structural disconnection was revealed as the least predictive measure. This suggests that detailed topographical information of voxel-wise data (represented via principal components) benefits model accuracy. This is in accordance with a previous investigation, where 2D pairwise disconnection outperformed 1D tract-wise disconnection (Griffis et al., 2019). In the present study, we used the same patient sample and a very similar prediction algorithm as in an earlier investigation (Röhrig et al., 2022). Thus, we were in the position to compare prediction performances of models using voxel-wise structural disconnections (present study) with voxel-wise anatomical maps (Röhrig et al., 2022). In sum, voxel-wise structural disconnections derived from stroke lesions alone achieved mean predictions as accurate as voxel-wise anatomical maps of stroke lesion and bilateral or right hemispheric WMH combined, explaining 42% of the total variance (cf. Röhrig et al., 2022). To this end, it is worth investing in high-dimensional measures, but the inclusion of WMH might not necessarily increase model performance. The estimation of individual stroke lesion-derived structural disconnectomes and the delineation and preprocessing of individual WMH topologies appear to be equally beneficial. We further demonstrated that stroke lesion-derived structural disconnections (voxel-wise 42% and pairwise 40% explained variance) outperformed voxel-wise stroke lesion anatomy (37% explained variance, cf. Röhrig et al., 2022), supporting previous studies that reported predictive superiority of structural disconnection measures over lesion anatomy (Griffis et al., 2019; Khalilian et al., 2024; Thiebaut de Schotten et al., 2020). In contrast, other studies reported similar prediction performances for anatomical and structural disconnection variables (Hope et al., 2018; Salvalaggio et al., 2020). It should therefore be explored in future studies under which specific circumstances structural disconnections can broaden insights into mechanisms of brain pathology beyond lesion anatomy itself.

To summarize, prediction performance can be similarly improved by either estimating structural disconnections caused by individual stroke lesions or by analyzing WMH anatomy alongside stroke anatomy, compared to using stroke anatomy alone (cf. Röhrig et al., 2022). Either way, voxel-wise data (represented via principal components) seem to achieve most accurate predictions. Future research should investigate whether features of lesion anatomy and structural disconnection (and functional disconnection) together outperform models that use only one measure of brain damage. Feature selection may be a promising option to identify the most predictive variable combination. However, some research already indicated that lesion-induced disconnectivity might be unable to give further information in addition to lesion anatomy, since the former is dependent on the latter (Halai et al., 2020; Hope et al., 2018; Zhao et al., 2023). On the other hand, a very recent investigation revealed that the combination of structural and functional disconnectivity (and demographics) achieved more accurate predictions than models with either one of them (and demographics) (Petersen et al., 2024).

### Conclusions

White matter hyperintensities (WMH) combined with the individual stroke lesion predominantly resulted in the disruption of fronto-subcortical pathways associated with spatial neglect. WMH appear to increase structural disconnections primarily in regions of the MFG, basal ganglia, and thalamus, with damage to the fronto-pontine tract being particularly critical. However, many neglect-related associations were less strong when WMH were considered, indicating that WMH may introduce noise. Future studies thus may investigate crucial characteristics of neglect-related WMH, as location and intensity, to reduce WMH-based noise in future lesion-mapping and prediction studies. Predictive modeling in the present study revealed that detailed topographical information of voxel-wise disconnection data seem to benefit model accuracy. Prediction performance could be similarly improved by either estimating structural disconnections caused by individual stroke lesions or by analyzing WMH anatomy alongside stroke anatomy, compared to using stroke anatomy alone. In conclusion, the structural disconnectome of premorbid WMH contributes to the understanding of neural correlates of spatial neglect; this kind of data may, however, not be predictive beyond the stroke-based disconnectome.

## Abbreviations

CoC: Center of Cancellation
GLM: general linear model
FWE: family-wise error
IFG: inferior frontal gyrus
IFJ: inferior frontal junction
IPL: inferior parietal lobule
MFG: middle frontal gyrus
MNI: Montreal Neurological Institute
MTG: middle temporal gyrus
STG: superior temporal gyrus
WMH: white matter hyperintensities

## Data availability

Online materials including overlay maps and result files are available online at Mendeley Data (DOI: 10.17632/fx3w28mzcv.1; https://data.mendeley.com/datasets/fx3w28mzcv/1). Patient-related data are not openly available due to data restriction and privacy guidelines of the local ethics commission of Tübingen University Clinic.

