## Supplementary Material for "Structural disconnections caused by white matter hyperintensities in post-stroke spatial neglect"

### Supplementary Analyses

#### Analysis of bilateral white matter hyperintensities

A recent simulation study by Röhrig et al. (2023) found that for unilateral brain damage (either in the left or right hemisphere), structural disconnections should be analyzed separately for each hemisphere. Combining them in a joint analysis can lead to a statistical imbalance, which disadvantages intra-hemispheric disconnections (Röhrig et al., 2023). In present analyses, we investigated patients with unilateral stroke damage and bilateral white matter hyperintensities (WMH). Since bilateral WMH affect both hemispheres simultaneously (despite some asymmetries), each type of disconnection can potentially occur in each individual (i.e. inter-hemispheric disconnections just as left- and right-sided intra-hemispheric disconnections). This circumstance results in inter- and intra-hemispheric disconnections being balanced when jointly analyzing left and right hemispheric WMH, indicating the unnecessity for separating bilateral WMH-related structural disconnections by hemisphere. For this reason, we here present joint analyses of bilateral WMH. Analyses using only right-sided WMH, however, are presented within the main manuscript; it was previously shown that left-sided WMH were less informative than right-sided WMH, and right-sided and bilateral WMH were almost equally predictive (Röhrig et al., 2022).

##### Voxel-wise results

When analyzing voxel-wise disconnectivity derived from the joint topography of stroke lesion and bilateral WMH, 221 voxels were significantly associated with neglect severity ($z > 5.93$). The voxel with the highest $z$-score of 6.84 is part of the right inferior frontal junction within the MFG (MNI-coordinates = 39, 8, 47). Among the white matter tracts, significant voxels were most frequently part of the right fronto-pontine tract, right cortico-striatal pathway, and right U-fibers (HCP-842; Yeh et al., 2018); significant voxels further cover white matter underlying the right SFG, MFG, and basal ganglia (BNA; Fan et al., 2016). We found 77 voxels to be more strongly associated to neglect severity when bilateral WMH were considered; the voxel with the largest increase is located at the right cortico-thalamic pathway next to the dorsal caudate (MNI-coordinates = 20, 2, 20).

#### Pairwise results

When combining stroke lesion and bilateral WMH, we identified significant disconnections: $N = 62$ at $p < 0.05$ ($t \geq 4.59$); $N = 14$ at $p < 0.01$ ($t \geq 5.38$); $N = 5$ at $p < 0.005$ ($t \geq 5.72$). In total, 31 gray matter regions were affected. The right globus pallidus was the only gray matter region with at least 10 disconnections. The most significant disconnections were found within the right hemisphere between the ventrolateral area 6 of the MFG and globus pallidus respectively dorsolateral putamen respectively posterior parietal thalamus, between globus pallidus and inferior frontal junction of the MFG, and between the dorsolateral area 8 of the SFG and dorsal area 9/46 of the MFG. We found 41 pairwise connections with a stronger association to neglect behavior when bilateral WMH were involved; the pairwise connection with the largest increase connects the right ventrolateral area 8 of the MFG and the right posterior parietal thalamus.

#### Tract-wise results

The joint topography of stroke lesion and bilateral WMH did reveal tracts significantly related to neglect severity: $N = 11$ at $p < 0.05$ ($t > 3.44$); $N = 7$ at $p < 0.01$ ($t > 4.29$); $N = 5$ at $p < 0.005$ ($t > 4.63$); $N = 3$ at $p < 0.001$ ($t > 5.34$). Among most significant tracts, we found the right U-fibers, right cortico-striatal pathway, and right fronto-pontine tract to be highly associated with neglect severity. We found 4 white matter tracts with a stronger association to neglect behavior when bilateral WMH were considered; the right fronto-pontine tract received the largest increase in association strength.

#### Parcel-wise results

When bilateral WMH were combined with the individual stroke lesion, findings revealed gray matter parcels significantly related to neglect severity: $N = 9$ at $p < 0.05$ ($t > 4.76$); $N = 3$ at $p < 0.01$ ($t > 5.52$); $N = 1$ at $p < 0.005$ ($t > 5.87$). The most significant parcel was rostral area 21 of the right MTG. We found no parcel that was stronger associated to neglect behavior when bilateral WMH were considered.

### Supplementary Tables

**Table S1. Number of features used for statistical analyses.**

| Disconnection origin | Disconnection type | | | |
|---|---|---|---|---|
| | Voxel-wise | Pairwise | Tract-wise | Parcel-wise |
| Stroke lesion | 330,557 | 2024 | 30 | 80 |
| Right-sided WMH | 305,699 | 2065 | 27 | 39 |
| Bilateral WMH | 483,673 | 2818 | 49 | 83 |
| Stroke lesion + right-sided WMH | 389,934 | 2177 | 31 | 102 |
| Stroke lesion + bilateral WMH | 559,905 | 2919 | 52 | 145 |

*Note.* To ignore features that were not at all or less frequently damaged (i.e. features with no or little variance), we restricted our main analyses (GLMs and prediction analyses) to features that were affected in a minimum number of patients. For voxel-wise data, we ignored voxels damaged in less than 5 patients, whereas for pairwise, tract-wise, and parcel-wise data, we removed features damaged in less than 10 patients. Total number of features before restriction: 985,860 voxels, 60,516 pairwise connections, 70 white matter tracts, and 246 gray matter parcels. Principal component analysis was performed on voxel-wise data after feature restriction for predictive modeling: for stroke lesion-induced disconnectivity, we kept 76 components cumulatively explaining > 99.5 % of the imaging variance; for right-sided WMH-induced disconnectivity, we kept 38 components cumulatively explaining > 90 % of the imaging variance; for bilateral WMH-induced disconnectivity, we kept 44 components cumulatively explaining > 90 % of the imaging variance; for stroke lesion and right-sided WMH-induced disconnectivity, we kept 39 components explaining > 90 % variance; for stroke lesion and bilateral WMH-induced disconnectivity, we kept 96 components explaining > 99.5 % variance (i.e. model-specific optima).

**Table S2. Most significant results of pairwise analyses.**

| Stroke lesion | | | Stroke lesion + right WMH | | | Stroke lesion + bilateral WMH | | |
|---|---|---|---|---|---|---|---|---|
| *Region A*<br>*Region B* | *t* | *p(FWE)* | *Region A*<br>*Region B* | *t* | *p(FWE)* | *Region A*<br>*Region B* | *t* | *p(FWE)* |
| MFG_R_A6vl<br>IFG_L_A44op | 7.123 | < 0.001 | MFG_ R_A6vl<br>BG_R_GP | 6.005 | < 0.001 | MFG_ R_A6vl<br>BG_R_GP | 6.005 | < 0.005 |
| MFG_R_A8vl<br>IFG_L_A44op | 6.698 | < 0.001 | MFG_R_IFJ<br>BG_R_GP | 5.990 | < 0.001 | MFG_R_IFJ<br>BG_R_GP | 5.990 | < 0.005 |
| MFG_R_IFJ<br>Tha_R_PPtha | 6.479 | < 0.005 | MFG_ R_A6vl<br>BG_R_dlPu | 5.856 | < 0.005 | MFG_ R_A6vl<br>BG_R_dlPu | 5.856 | < 0.005 |
| MFG_R_A8vl<br>INS_L_dIa | 6.453 | < 0.005 | MFG_ R_A6vl<br>Tha_R_PPtha | 5.851 | < 0.005 | MFG_ R_A6vl<br>Tha_R_PPtha | 5.851 | < 0.005 |
| MFG_ R_A8vl<br>IFG_L_A44v | 6.451 | < 0.005 | SFG_R_A8dl<br>MFG_R_A9/46d | 5.735 | < 0.005 | SFG_R_A8dl<br>MFG_R_A9/46d | 5.735 | < 0.005 |
| MFG_R_IFJ<br>BG_R_dCa | 6.372 | < 0.005 | MFG_R_IFJ<br>BG_R_dlPu | 5.715 | < 0.005 | MFG_R_IFJ<br>BG_R_dlPu | 5.715 | < 0.01 |
| MFG_ R_A6vl<br>IFG_L_A44v | 6.309 | < 0.005 | MFG_ R_A6vl<br>Tha_R_cTtha | 5.632 | < 0.005 | MFG_ R_A6vl<br>Tha_R_cTtha | 5.632 | < 0.01 |
| MFG_L_A8vl<br>MFG_ R_A6vl | 6.265 | < 0.005 | MFG_ R_A6vl<br>Tha_R_mPMtha | 5.626 | < 0.005 | MFG_ R_A6vl<br>Tha_R_mPMtha | 5.626 | < 0.01 |
| MFG_R_IFJ<br>IPL_R_A39rd | 6.179 | < 0.005 | SFG_R_A8dl<br>BG_R_GP | 5.565 | < 0.005 | SFG_R_A8dl<br>BG_R_GP | 5.565 | < 0.01 |
| MFG_L_A6vl<br>MFG_ R_A6vl | 6.088 | < 0.005 | MFG_ R_A6vl<br>Tha_R_lPFtha | 5.541 | < 0.005 | MFG_ R_A6vl<br>Tha_R_lPFtha | 5.541 | < 0.01 |
| MFG_ R_A6vl<br>INS_L_vIa | 6.034 | < 0.005 | MFG_R_IFJ<br>Tha_R_mPMtha | 5.502 | < 0.005 | MFG_R_IFJ<br>Tha_R_mPMtha | 5.502 | < 0.01 |
| MFG_R_A9/46d<br>MFG_L_A8vl | 5.966 | < 0.005 | SFG_R_A6dl<br>BG_R_dlPu | 5.416 | < 0.005 | SFG_R_A6dl<br>BG_R_dlPu | 5.416 | < 0.01 |
| STG_R_A38l<br>IPL_L_A39rd | 5.946 | < 0.005 | MTG_L_A21r<br>PhG_R_TI | 5.414 | < 0.005 | STG_R_A38m<br>INS_R_vIa | 5.398 | < 0.01 |
| MTG_R_A21r<br>LOcC_R_msOccG | 5.879 | < 0.005 | STG_R_A38m<br>INS_R_vIa | 5.398 | < 0.005 | SFG_R_A8dl<br>Tha_R_PPtha | 5.393 | < 0.01 |
| MFG_ R_A8vl<br>INS_L_dIg | 5.877 | < 0.005 | SFG_R_A8dl<br>Tha_R_PPtha | 5.393 | < 0.005 | MFG_ R_A8vl<br>BG_R_GP | 5.356 | < 0.05 |
| MFG_R_IFJ<br>MFG_L_A6vl | 5.874 | < 0.005 | MFG_ R_A8vl<br>BG_R_GP | 5.356 | < 0.005 | IFG_R_A44d<br>Tha_R_lPFtha | 5.278 | < 0.05 |
| MFG_ R_A6vl<br>INS_L_dIa | 5.868 | < 0.005 | IFG_R_A44d<br>Tha_R_lPFtha | 5.278 | < 0.005 | MFG_ R_A8vl<br>Tha_R_mPMtha | 5.220 | < 0.05 |
| MFG_ R_A6vl<br>BG_R_dCa | 5.857 | < 0.005 | MFG_ R_A6vl<br>IFG_L_A44v | 5.275 | < 0.005 | SFG_R_A8dl<br>Tha_R_lPFtha | 5.194 | < 0.05 |
| STG_R_A38l<br>MTG_L_A21r | 5.855 | < 0.005 | MFG_ R_A8vl<br>Tha_R_mPMtha | 5.220 | < 0.01 | IFG_R_A44d<br>Tha_R_mPMtha | 5.193 | < 0.05 |
| STG_R_A38l<br>ITG_L_A20cl | 5.852 | < 0.005 | SFG_R_A8dl<br>Tha_R_lPFtha | 5.194 | < 0.01 | SFG_R_A6m<br>BG_R_dlPu | 5.192 | < 0.05 |
| STG_R_A38l<br>ITG_L_A37elv | 5.839 | < 0.005 | IFG_R_A44d<br>Tha_R_mPMtha | 5.193 | < 0.01 | SFG_R_A8dl<br>Tha_R_mPMtha | 5.190 | < 0.05 |
| MFG_R_IFJ<br>BG_R_dlPu | 5.834 | < 0.005 | SFG_R_A6m<br>BG_R_dlPu | 5.192 | < 0.01 | SFG_R_A8m<br>BG_R_dlPu | 5.170 | < 0.05 |
| MTG_L_A21r<br>MTG_R_A21r | 5.831 | < 0.005 | SFG_R_A8dl<br>Tha_R_mPMtha | 5.190 | < 0.01 | MFG_ R_A6vl<br>Tha_R_Otha | 5.164 | < 0.05 |
| MFG_R_IFJ<br>Tha_R_lPFtha | 5.805 | < 0.005 | SFG_R_A8m<br>BG_R_dlPu | 5.170 | < 0.01 | SFG_R_A8dl<br>BG_R_dlPu | 5.163 | < 0.05 |
| STG_R_A38l<br>LOcC_L_V5/MT+ | 5.795 | < 0.005 | MFG_ R_A6vl<br>Tha_R_Otha | 5.164 | < 0.01 | MFG_ R_A8vl<br>Tha_R_PPtha | 5.082 | < 0.05 |

*Note*. Table reports 25 region pairs most significantly associated to neglect severity, sorted by decreasing *t*-statistics. Results were obtained from GLMs using pairwise structural disconnection data. *P*-thresholds were obtained using permutation-based FWE-correction. Disconnections were derived by either stroke lesions or stroke lesions and right-sided WMH combined. Total number of significant pairwise disconnections: *N(stroke lesion)* = 394, *N(stroke lesion and right WMH)* = 112, *N(stroke lesion and bilateral WMH)* = 62. Parcels and respective abbreviations are based on the Brainnetome Atlas with 246 regions (Fan et al., 2016).

*Abbreviations*: BG – basal ganglia, IFG – inferior frontal gyrus, INS – insular gyrus, IPL – inferior parietal lobe, ITG – inferior temporal gyrus, L – left hemisphere, LOcC – lateral occipital cortex, MFG – middle frontal gyrus, MTG – middle temporal gyrus, PhG – parahippocampal gyrus, R – right hemisphere, SFG – superior frontal gyrus, STG – superior temporal gyrus, Tha - thalamus.

**Table S3. Region counts of pairwise disconnections.**

| Stroke lesion | | Stroke lesion + right WMH | | Stroke lesion + bilateral WMH | |
|---|---|---|---|---|---|
| *Region* | *Count* | *Region* | *Count* | *Region* | *Count* |
| STG_R_A38l | 28 | MFG_R_A6vl | 15 | BG_R_GP | 10 |
| MFG_R_A6vl | 26 | BG_R_GP | 15 | | |
| ITG_R_A37vl | 19 | BG_R_dlPu | 15 | | |
| MFG_R_IFJ | 18 | SFG_R_A8dl | 11 | | |
| MFG_R_A8vl | 18 | MFG_R_A8vl | 11 | | |
| PrG_R_A4tl | 18 | Tha_R_mPMtha | 10 | | |
| MTG_R_A37dl | 18 | | | | |
| IFG_R_A44v | 17 | | | | |
| PrG_R_A4hf | 16 | | | | |
| ITG_R_A37elv | 16 | | | | |
| ITG_R_A20iv | 15 | | | | |
| ITG_R_A20cl | 15 | | | | |
| MTG_R_A21r | 14 | | | | |
| IFG_R_A44d | 12 | | | | |
| MTG_L_A21r | 12 | | | | |
| INS_R_dId | 12 | | | | |
| BG_R_GP | 12 | | | | |
| BG_R_dlPu | 12 | | | | |
| SFG_R_A8dl | 11 | | | | |
| IPL_L_A39c | 11 | | | | |
| IPL_R_A39rd | 11 | | | | |
| IPL_R_A40rd | 11 | | | | |

*Note*. Table reports all regions that were involved in at least 10 pairwise structural disconnections significantly associated to neglect severity ($p$ < 0.05, with permutation-based FWE-correction). Disconnections were derived by either stroke lesions or stroke lesions and right-sided WMH combined. Parcels and respective abbreviations are based on the Brainnetome Atlas with 246 regions (Fan et al., 2016). For visualization, see Figure S1.

*Abbreviations*: BG – basal ganglia, IFG – inferior frontal gyrus, INS – insular gyrus, IPL – inferior parietal lobe, ITG – inferior temporal gyrus, L – left hemisphere, MFG – middle frontal gyrus, MTG – middle temporal gyrus, PrG – precentral gyrus, R – right hemisphere, SFG – superior frontal gyrus, STG – superior temporal gyrus, Tha - thalamus.

**Table S4. Significant results of tract-wise analyses.**

| Stroke lesion | | | Stroke lesion + right WMH | | | Stroke lesion + bilateral WMH | | |
|---|---|---|---|---|---|---|---|---|
| *Tract* | *t* | *p(FWE)* | *Tract* | *t* | *p(FWE)* | *Tract* | *t* | *p(FWE)* |
| U_R | 7.551 | < 0.001 | U_R | 5.973 | < 0.001 | U_R | 5.973 | < 0.001 |
| SLF_R | 6.158 | < 0.001 | CS_R | 5.358 | < 0.001 | CS_R | 5.358 | < 0.001 |
| CT_R | 5.823 | < 0.001 | FPT_R | 5.339 | < 0.001 | FPT_R | 5.339 | < 0.001 |
| CCMidAnterior | 5.597 | < 0.001 | CT_R | 5.174 | < 0.001 | CT_R | 5.174 | < 0.005 |
| CS_R | 5.467 | < 0.001 | SLF_R | 4.750 | < 0.005 | SLF_R | 4.750 | < 0.005 |
| CCCentral | 5.204 | < 0.001 | UF_R | 4.423 | < 0.005 | UF_R | 4.423 | < 0.01 |
| AF_R | 4.963 | < 0.005 | CST_R | 4.346 | < 0.005 | CST_R | 4.346 | < 0.01 |
| FPT_R | 4.889 | < 0.005 | PPT_R | 4.192 | < 0.005 | PPT_R | 4.192 | < 0.05 |
| CCPosterior | 4.880 | < 0.005 | AC | 4.152 | < 0.01 | AC | 4.168 | < 0.05 |
| CCAnterior | 4.876 | < 0.005 | CCMidAnterior | 4.050 | < 0.01 | FAT_R | 4.011 | < 0.05 |
| FAT_R | 4.594 | < 0.005 | FAT_R | 4.011 | < 0.01 | EMC_R | 3.718 | < 0.05 |
| EMC_R | 4.422 | < 0.005 | EMC_R | 3.718 | < 0.05 | | | |
| CCMidPosterior | 4.359 | < 0.005 | CCCentral | 3.424 | < 0.05 | | | |
| UF_R | 4.255 | < 0.01 | AF_R | 3.253 | < 0.05 | | | |
| AR_R | 4.239 | < 0.01 | | | | | | |
| AC | 4.231 | < 0.01 | | | | | | |
| CST_R | 4.128 | < 0.01 | | | | | | |
| TPT_R | 3.996 | < 0.05 | | | | | | |
| PPT_R | 3.915 | < 0.05 | | | | | | |
| OPT_R | 3.874 | < 0.05 | | | | | | |
| MdLF_R | 3.630 | < 0.05 | | | | | | |
| IFOF_R | 3.313 | < 0.05 | | | | | | |

*Note*. Table reports all white matter tracts that were significantly associated to neglect severity, sorted by decreasing *t*-statistics. Results were obtained from GLMs using tract-wise structural disconnection data. Disconnections were derived by either stroke lesions or stroke lesions and right-sided WMH combined. *P*-thresholds were obtained using permutation-based FWE-correction. Tracts and respective abbreviations are based on the HCP-842 tractography atlas (Yeh et al., 2018) and the segmentation of the corpus callosum (FreeSurferSeg ROIs distributed with DSI Studio, https://dsi-studio.labsolver.org/) with total 70 tracts.

*Abbreviations*: AC – anterior commissure, AF – arcuate fasciculus, AR – acoustic radiation, CC – corpus callosum, CS – cortico-striatal pathway, CST – cortico-spinal tract, CT – cortico-thalamic pathway, EMC – extreme capsule, FAT – frontal-aslant tract, FPT – fronto-pontine tract, IFOF – inferior fronto-occipital fasciculus, MdLF – middle longitudinal fasciculus, OPT – occipito-pontine tract, PPT – parieto-pontine tract, R – right hemisphere, SLF – superior longitudinal fasciculus, TPT- temporo-pontine tract, U – U-fiber, UF – uncinate fasciculus.

**Table S5. Significant results of parcel-wise analyses.**

| Stroke lesion | | | Stroke lesion + right WMH | | | Stroke lesion + bilateral WMH | | |
|---|---|---|---|---|---|---|---|---|
| *Parcel* | *t* | *p(FWE)* | *Parcel* | *t* | *p(FWE)* | *Parcel* | *t* | *p(FWE)* |
| MTG_A21r | 6.198 | < 0.005 | MTG_A21r | 6.197 | < 0.005 | MTG_A21r | 6.197 | < 0.005 |
| STG_A38m | 5.752 | < 0.005 | STG_A38m | 5.752 | < 0.005 | STG_A38m | 5.752 | < 0.01 |
| PrG_A6cvl | 5.628 | < 0.005 | PrG_A6cvl | 5.608 | < 0.01 | PrG_A6cvl | 5.608 | < 0.01 |
| STG_A38l | 5.313 | < 0.01 | STG_A38l | 5.311 | < 0.01 | STG_A38l | 5.311 | < 0.05 |
| MFG_IFJ | 5.288 | < 0.01 | MFG_IFJ | 5.284 | < 0.05 | MFG_IFJ | 5.284 | < 0.05 |
| PrG_A4tl | 5.226 | < 0.05 | PrG_A4tl | 5.222 | < 0.05 | PrG_A4tl | 5.222 | < 0.05 |
| MFG_A6vl | 5.063 | < 0.05 | MFG_A6vl | 5.035 | < 0.05 | MFG_A6vl | 5.035 | < 0.05 |
| INS_vId/vIg | 4.954 | < 0.05 | INS_vId/vIg | 4.954 | < 0.05 | INS_vId/vIg | 4.954 | < 0.05 |
| IFG_A44v | 4.798 | < 0.05 | IFG_A44v | 4.791 | < 0.05 | IFG_A44v | 4.791 | < 0.05 |
| IPL_A39c | 4.728 | < 0.05 | IPL_A39c | 4.738 | < 0.05 | | | |
| STG_TE1.0/.2 | 4.492 | < 0.05 | | | | | | |
| OrG_A11l | 4.445 | < 0.05 | | | | | | |
| BG_GP | 4.404 | < 0.05 | | | | | | |

*Note*. Table reports all parcels that were significantly associated to neglect severity, sorted by decreasing *t*-statistics. Results were obtained from GLMs using parcel-wise structural disconnection data. Disconnections were derived by either stroke lesions or stroke lesions and right-sided WMH combined. *P*-thresholds were obtained using permutation-based FWE-correction. Parcels and respective abbreviations are based on the Brainnetome Atlas with 246 regions (Fan et al., 2016). All reported parcels are right hemispheric. For visualization, see Figure S2.

*Abbreviations*: BG – basal ganglia, IFG – inferior frontal gyrus, INS – insular gyrus, IPL – inferior parietal lobe, MFG – middle frontal gyrus, MTG – middle temporal gyrus, OrG – orbital gyrus, PrG – precentral gyrus, STG – superior temporal gyrus.

**Table S6. WMH-related increase in statistical association strength.**

| Pairwise | | | | Tract-wise | | | |
|---|---|---|---|---|---|---|---|
| *Region A*<br>*Region B* | *t*<br>*(stroke)* | *t (stroke*<br>*+ WMH)* | Δ*t* | *Tract* | *t*<br>*(stroke)* | *t (stroke*<br>*+ WMH)* | Δ*t* |
| MFG_R_A8vl<br>Tha_R_PPtha | 4.209 | 5.082 | 0.873 | FPT_R | 4.889 | 5.339 | 0.450 |
| PrG_R_A4t<br>BG_R_GP | 3.641 | 4.489 | 0.849 | PPT_R | 3.915 | 4.192 | 0.277 |
| PCL_R_A4ll<br>BG_R_dlPu | 3.575 | 4.421 | 0.847 | CST_R | 4.128 | 4.346 | 0.217 |
| SFG_R_A8dl<br>Tha_R_mPMtha | 4.348 | 5.190 | 0.842 | UF_R | 4.255 | 4.423 | 0.168 |
| SFG_R_A8dl<br>PrG_R_A6cdl | 3.432 | 4.224 | 0.793 | | | | |
| MFG_R_A9/46d<br>Tha_R_rTtha | 4.254 | 5.014 | 0.760 | | | | |
| SFG_R_A8dl<br>BG_R_GP | 4.828 | 5.565 | 0.736 | | | | |
| SFG_R_A6dl<br>Tha_R_mPMtha | 4.109 | 4.832 | 0.723 | | | | |
| IFG_R_A44d<br>Tha_R_lPFtha | 4.564 | 5.278 | 0.714 | | | | |
| SFG_R_A8m<br>Tha_R_mPMtha | 4.091 | 4.799 | 0.708 | | | | |

*Note*. Table reports pairwise connections and white matter tracts with a stronger association to neglect severity when right-sided WMH were considered. For pairwise results, top 10 connections with the largest increase are presented (from a total of 56 connections). To identify connections/tracts that were more strongly associated with neglect severity due to WMH involvement, we only considered connections/tracts that were tested in both analyses (stroke lesion with and without WMH), that resulted in $\Delta t > 0.01$, and that reached significance when WMH were considered ($p < 0.05$). Parcels and respective abbreviations are based on the Brainnetome Atlas with 246 regions (Fan et al., 2016). Tracts and respective abbreviations are based on the HCP-842 tractography atlas (Yeh et al., 2018) and the segmentation of the corpus callosum (FreeSurferSeg ROIs distributed with DSI Studio, https://dsi-studio.labsolver.org/) with total 70 tracts.

*Abbreviations*: BG – basal ganglia, CST – cortico-spinal tract, FPT – fronto-pontine tract, IFG – inferior frontal gyrus, MFG – middle frontal gyrus, PCL – paracentral lobule, PPT – parieto-pontine tract, PrG – precentral gyrus, R – right hemispheric, SFG – superior frontal gyrus, Tha – thalamus, UF – uncinate fasciculus.

#### Supplementary Figures

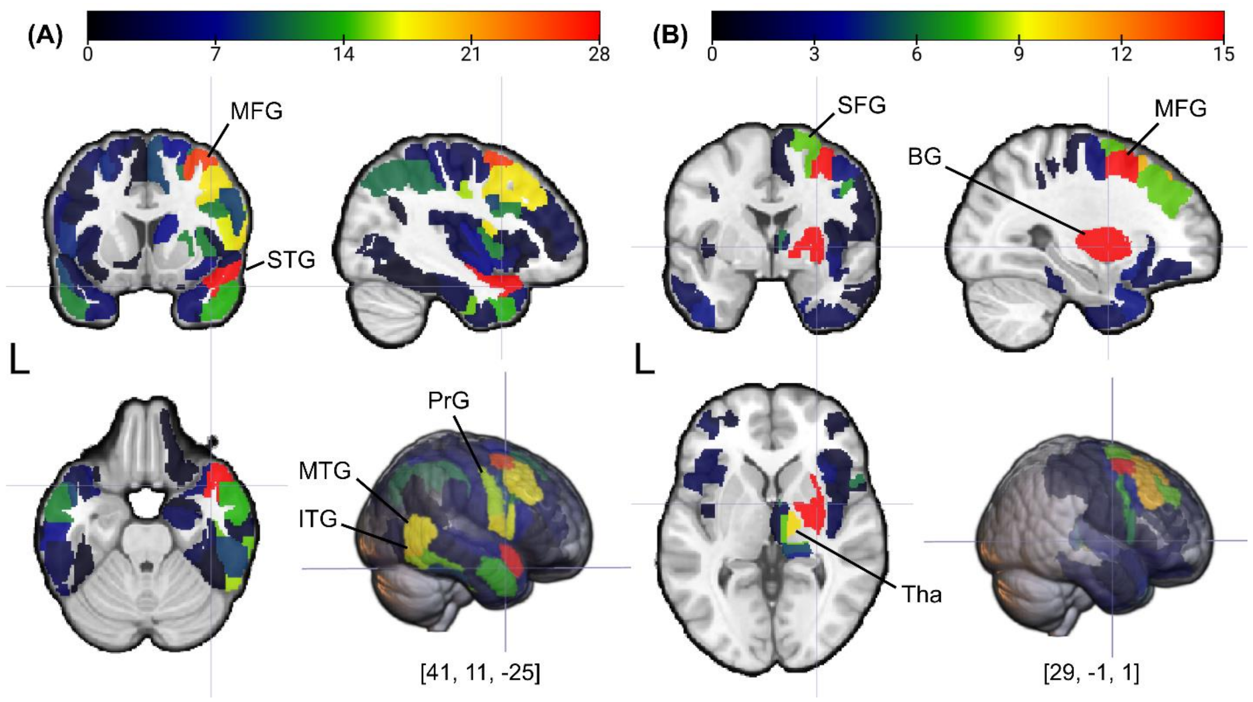


**Figure S1. Brain regions most frequently involved in significant pairwise structural disconnections.** Cortical and subcortical brain parcels of the Brainnetome Atlas (Yeh et al., 2018) are color-coded based on respective disconnection frequencies, i.e. how often a region was involved in region-to-region disconnections significantly associated to neglect severity ($p < 0.05$, with permutation-based FWE-correction). Pairwise disconnections were derived by either **(A)** stroke lesions or **(B)** stroke lesions and right-sided WMH combined. Coronal, sagittal, and axial views as well as the whole brain are presented; crosshair is located at (one of) the most frequently disconnected region (MNI coordinates are reported in square brackets). For details, see Table S3.

*Abbreviations*: BG – basal ganglia, ITG – inferior temporal gyrus, L – left hemisphere, MFG – middle frontal gyrus, MTG – middle temporal gyrus, PrG – precentral gyrus, SFG – superior frontal gyrus, STG – superior temporal gyrus, Tha - thalamus.

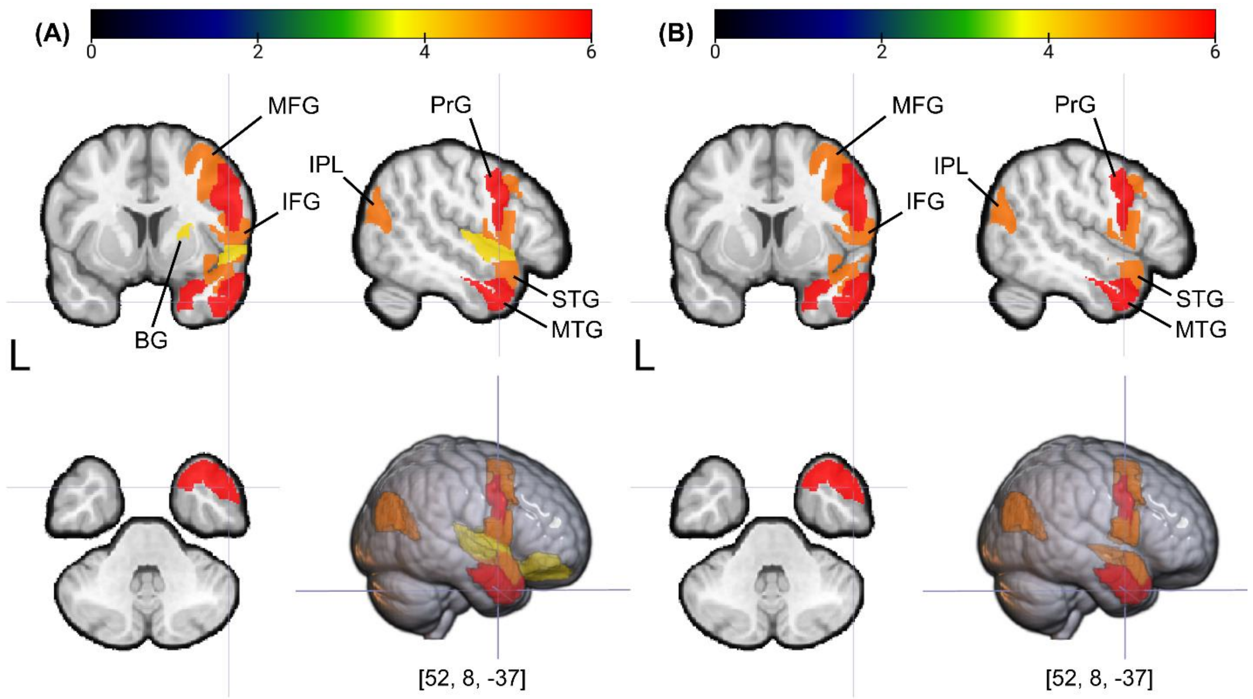


**Figure S2. Parcel-wise results.** Significant cortical and subcortical brain parcels of the Brainnetome Atlas (Yeh et al., 2018) are color-coded based on respective *t*-statistics, i.e. how strongly parcel-wise structural disconnection was associated to neglect severity ($p < 0.05$, with permutation-based FWE-correction). Parcel-wise disconnections were derived by either **(A)** stroke lesions or **(B)** stroke lesions and right-sided WMH combined. Coronal, sagittal, and axial views as well as the whole brain are presented; crosshair is located at the most strongly associated region (MNI coordinates are reported in square brackets). For details, see Table S5.

*Abbreviations*: BG – basal ganglia, IFG – inferior frontal gyrus, IPL – inferior parietal lobe, L – left hemisphere, MFG – middle frontal gyrus, MTG – middle temporal gyrus, PrG – precentral gyrus, STG – superior temporal gyrus.
